# Marker-based estimates reveal significant non-additive effects in clonally propagated cassava (*Manihot esculenta*): implications for the prediction of total genetic value and the selection of varieties

**DOI:** 10.1101/031864

**Authors:** Marnin D. Wolfe, Peter Kulakow, Ismail Y. Rabbi, Jean-Luc Jannink

## Abstract

In clonally propagated crops, non-additive genetic effects can be effectively exploited by the identification of superior genetic individuals as varieties. Cassava *(Manihot esculenta* Crantz) is a clonally propagated staple food crop that feeds hundreds of millions. We quantified the amount and nature of non-additive genetic variation for key traits in a breeding population of cassava from sub-Saharan Africa using additive and non-additive genome-wide marker-based relationship matrices. We then assessed the accuracy of genomic prediction of additive compared to total (additive plus non-additive) genetic value. We confirmed previous findings based on diallel populations, that non-additive genetic variation is significant, especially for yield traits. Further, we show that we total genetic value correlated more strongly to observed phenotypes than did additive value, although this is constrained by low broad-sense heritability and is not beneficial for traits with already high heritability. We address the implication of these results for cassava breeding and put our work in the context of previous results in cassava, and other plant and animal species.

## INTRODUCTION

Understanding genetic architecture requires the decomposition of genetic effects into additive, dominance, and epistatic components (Fisher 1918; Cockerham 1954; Kempthorne 1954). However, partitioning genetic variance components is notoriously difficult, requiring specialized breeding designs (e.g. diallel crosses) and pedigree information (Lynch and Walsh 1998) often limiting the genetic diversity that can be sampled in any one given study. Genome-wide molecular marker data now enable the accurate measurement of relatedness in the form of genomic realized relationship matrices (GRMs) (VanRaden 2008; Heffner *et al*. 2009; Lorenz *et al*. 2011b). GRMs provide more accurate relatedness information than pedigrees because they directly measure Mendelian sampling (causing variation in relatedness within relatedness classes such as full-siblings) (Heffner *et al*. 2009). Further, GRMs can measure relationships even in diverse, nominally unrelated samples expanding the potential for studying inheritance in natural and breeding populations (Lorenz *et al*. 2011a).

Estimation of narrow-sense heritability and prediction of breeding values in genomic selection programs is becoming increasingly common using additive formulations of GRMs (Visscher *et al*. 2008). Several recent studies have described dominance and epistatic GRMs for the partitioning of non-additive genetic variance using genome-wide SNP markers (Su *et al*. 2012; Vitezica *et al*. 2013; Muñoz *et al*. 2014; Wang *et al*. 2014). Models using these new formulations have been shown to provide improved partitioning of genetic variances relative to pedigree-based approaches (Su *et al*. 2012; Muñoz *et al*. 2014). These new models can be used not only to estimate genetic variances but also for genomic prediction of additive and total genetic value in genomic selection breeding programs (Su *et al*. 2012; Vitezica *et al*. 2013; Muñoz *et al*. 2014; Wang *et al*. 2014).

Cassava is a vegetatively propagated, staple food crop that is high in starch and feeds half a billion people worldwide (http://faostat.fao.org). Efforts to improve cassava genetically with cutting edge methodologies including transgenic and genomic selection (GS) approaches are underway thanks to new genomic resources (Prochnik *et al*. 2012; (ICGMC) 2014). Prediction of additive genetic value has recently been evaluated (Oliveira *et al*. 2012; Ly *et al*. 2013) and genomic selection using standard models is currently being tested (http://www.nextgencassava.org). Vegetatively propagated crop (e.g. cassava) breeding can exploit non-additive genetic effects by identifying superior clones as varieties (Ceballos *et al*. 2015).

Diallel studies in cassava indicate that non-additive genetic effects (i.e. specific combining ability) are strong, particularly for root yield traits (Cach *et al*. 2005, 2006; Calle *et al*. 2005; Jaramillo *et al*. 2005; Perez *et al*. 2005; Pérez *et al*. 2005; Zacarias and Labuschagne 2010; Kulembeka *et al*. 2012; Tumuhimbise *et al*. 2014; Ceballos *et al*. 2015; Chalwe *et al*. 2015). If the limited number of parents tested thus far represents the broader cassava breeding germplasm, genetic gains, especially for already low-heritability root yield traits will be slow regardless of the breeding scheme employed (e.g. phenotypic vs. pedigree vs. genomic selection). Breeding gains have indeed been slow in cassava (Ceballos *et al*. 2012) and low accuracies have been reported for genomic prediction of yield compared with cassava mosaic disease (CMD) resistance and dry matter (DM) content (Oliveira *et al*. 2012; Ly *et al*. 2013). However, cassava varieties are evaluated and disseminated to farmers by clonal propagation, meaning that accurate prediction of total (additive plus non-additive) genetic value could contribute to variety selection.

In this study, we test the hypothesis that certain cassava traits, especially root yield have relatively large non-additive genetic variance that account for low genomic prediction accuracies previously observed. We estimate additive and non-additive variance components using genomic relationship matrices in two populations of cassava from the International Institute of Tropical Agriculture’s (IITA) genomic selection breeding program. Further, we assess the accuracy of predicting additive and total genetic value using the additive and non-additive models. We discuss the origin of non-additive genetic variance in cassava, its potential effect on cassava breeding, and its role in genomic selection strategies for cassava improvement in the future.

## METHODS

### Germplasm and Phenotyping Trials

We examined additive and non-additive effects in two populations of cassava that have been genotyped and phenotyped as part of the Next Generation Cassava Breeding Program at IITA, Nigeria (http://www.nextgencassava.org). The IITA’s Genetic Gain (GG) population contains 694 historically important clones, most of which are advanced breeding lines although some are classified as superior landraces. These lines have been selected and maintained clonally since 1970 (Okechukwu and Dixon 2008; Ly *et al*. 2013). Most of these materials are derived from the cassava gene pool from West Africa as well as parents derived from the breeding program at Amani Station in Tanzania and hybrids of germplasm introduced from Latin America. Available information on the GG accessions included in our analyses is provided in Table S1.

IITA’s Genetic Gain trials were conducted in seven locations over 14 years (2000 to 2014) in Nigeria for a total of 24,373 observations. Each GG trial comprises a randomized, incomplete block design replicated one or two times per location and year. Since materials have been occasionally lost and new, selected materials are continuously added to the GG, the number of clones trialed in a given year changes gradually across years, generally increasing. The sample sizes, number of replicates and number of clones from the GG in each of the trials (location-year combinations) is provided in Table S2.

Theory suggests that founding events and truncation selection can both lead to a conversion of non-additive genetic variation into additive variance. This can happen because of the induction of linkage disequilibrium and reduction in allele frequency (or fixation of alleles) at some loci relative to others (Goodnight 1988; Turelli and Barton 2006; Hallander and Waldmann 2007). Consequently, our results might depend on the population examined. We therefore analyzed an additional population: a collection of 2187 clones that are the direct descendants of truncation selection on the GG. Briefly, in 2012 the GG and all available historical phenotype data was used as a reference population to obtain genome estimated breeding values (GEBVs) using the genomic BLUP (GBLUP) model (VanRaden 2008; Heffner *et al*. 2009). Selection was based on an index that included mean cassava mosaic disease severity (MCMDS), mean cassava bacterial blight disease severity (MCBBS), dry matter content (DM), harvest index (HI) and fresh root weight (RTWT). This index of GEBVs was used to select 83 members of the GG to cross and generated a population of 137 full-sib families, which we refer to as the GS Cycle 1 (C1). The pedigree of the C1 is available in Table S3.

Cycle 1 progenies were evaluated in a single clonal evaluation trial during the 2013-2014 field season across three-locations (Ibadan, Ikenne, and Mokwa). For the C1 clonal trial, planting material was only available for one plot of five stands per clone, so each clone was only planted in one of the three locations (Table S2). Clones were assigned to each location so as to equally represent each family in every environment.

For both populations, we analyzed the five traits described above (MCMDS, MCBBS, DM, HI, RTWT) plus sprouting ability (SPROUT). MCMDS and MCBBS were scored on a scale of 1 (no symptoms) to 5 (severe symptoms). Most GG trials measured dry matter (DM) by the oven drying method although some trials used the specific gravity method. DM is expressed as a percentage of the fresh weight of roots. Fresh root weight (RTWT) is measured in kilograms per plot and harvest index (HI) is the percent of total biomass per plot (roots plus shoots) that is RTWT. Sprouting ability (SPROUT) was expressed as the percent of planted stakes sprouting at one month after planting.

### Genotype data

We used genotyping-by-sequencing (GBS) to obtain genome-wide SNP marker data (Elshire *et al*. 2011). We used the ApeKI restriction enzyme as recommended by (Hamblin and Rabbi 2014). SNPs were called using the TASSEL GBS pipeline V4 (Glaubitz *et al*. 2014) and aligned to the cassava reference genome, version 5, which is available on Phytozome (http://phytozome.jgi.doe.gov) and described by the International Cassava Genetic Map Consortium ((ICGMC) 2014). We removed individuals with more than 80% and markers with >60% missing genotype calls. Also, markers with extreme deviation from Hardy-Weinberg equilibrium (Chi-square > 20) were removed. If there were not at least two reads at a given locus for a given clone, the genotype was set to missing and imputed. SNP marker data was converted to the dosage format (0, 1, 2) and missing data were imputed with the glmnet algorithm in R (http://cran.r-project.org/web/packages/glmnet/index.html) as described in (Wong *et al*. 2014). We used 114,922 markers that passed these filters with a minor allele frequency greater than 1% to calculate genomic relationship matrices as described below.

### Genomic Relationship Matrices

We measured the realized additive, dominance and epistatic relationships in our population using functions that have been described previously (Su *et al*. 2012; Muñoz *et al*. 2014). We used the additive realized relationship matrix, A as described by Van Raden (VanRaden 2008): 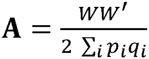. Here W is a matrix of dimension *n* individuals by *m* SNP markers. The elements of W are the marker dosages (0 – 2p_i_) for aa, (1 – 2p_i_) for Aa and (2 – 2p_i_) for AA, where p_i_ is the frequency of the second allele and q_i_ is the frequency of the first allele, at the *i*th locus. The a (or 0) allele refers to the reference genome allele. The **A** matrix was calculated using the *A.mat* function in the *rrBLUP* package (Endelman 2011).

As described by Su et al. (Su *et al*. 2012) the dominance relationship matrix, **D** is 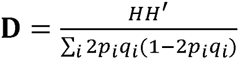 (also see (Muñoz *et al*. 2014)). Where H has the same dimensions as W, heterozygotes are scored (1 – 2p_i_q_i_) and homozygotes are (0 – 2p_i_q_i_). We made a custom modification (available upon request) to the *A.mat* function to produce the **D** matrix. Relationship matrices that capture epistasis can be calculated by taking the hadamard product (element-by-element multiplication; denoted #) of two or more matrices (Henderson 1985). For simplicity, we only explored additive-by-additive (**A#A**) and additive-by-dominance (**A#D**) relationships in this study.

### Variance component and heritability models

#### Single-step, Multi-environment

We used several approaches to estimate the relative importance of additive and non-additive effects in the Genetic Gain and Cycle 1 populations. In the first analysis, we analyzed the multi-year, multi-location GG data with a single random effects model. Since the entire historical phenotype dataset is large (24,373 observations) and was relatively unbalanced in sample size across years and locations, we only analyzed data from trials with >400 individuals. This filter resulted in a dataset of 7745 observations from three locations (Ibadan, Ubiaja, Mokwa) and eight years (2006-2014, excluding 2012). All 694 genotyped GG clones were represented in this dataset (Table S2).

The models we fit were similar to those described in Ly et al. (Ly *et al*. 2013). The full model was specified as follows: 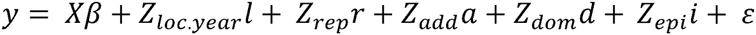. Here, y represents raw phenotypic observations. In our data, the only fixed effect (β) was an intercept for all traits except RTWT, which contained a covariate accounting for variation in the number of plants harvested per plot. The random effects terms for experimental design terms included a unique intercept for each trial (i.e. location-year combination), 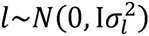, where I is the identity matrix and 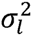 is the associated variance component as well as a replication effect, nested in location-year combination, 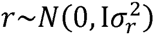.

The genetic variance component terms included 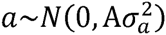, where A is the additive relationship matrix and 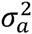 is the additive genetic variance component. Similarly, 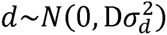, is the dominance effect with covariance D equal to the dominance relationship matrix and 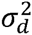 equal to the dominance variance. The epistatic term 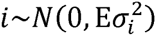 where the covariance matrix E took the form either of the A#A matrix (additive-by-additive) or the A#D matrix (additive-by-dominance) and the epistatic variance 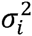 was correspondingly either 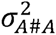or 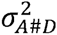. The final is the residual variance, assumed to be random and distributed 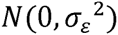. The terms X, Z_loc.year_, Z_rep_, Z_add_, Z_dom_ and Z_epi_ are incidence matrices relating observations to the levels of each factor. We list the different models fit in Table 1, each of which are variations on the full model described above.

**Table 1.**
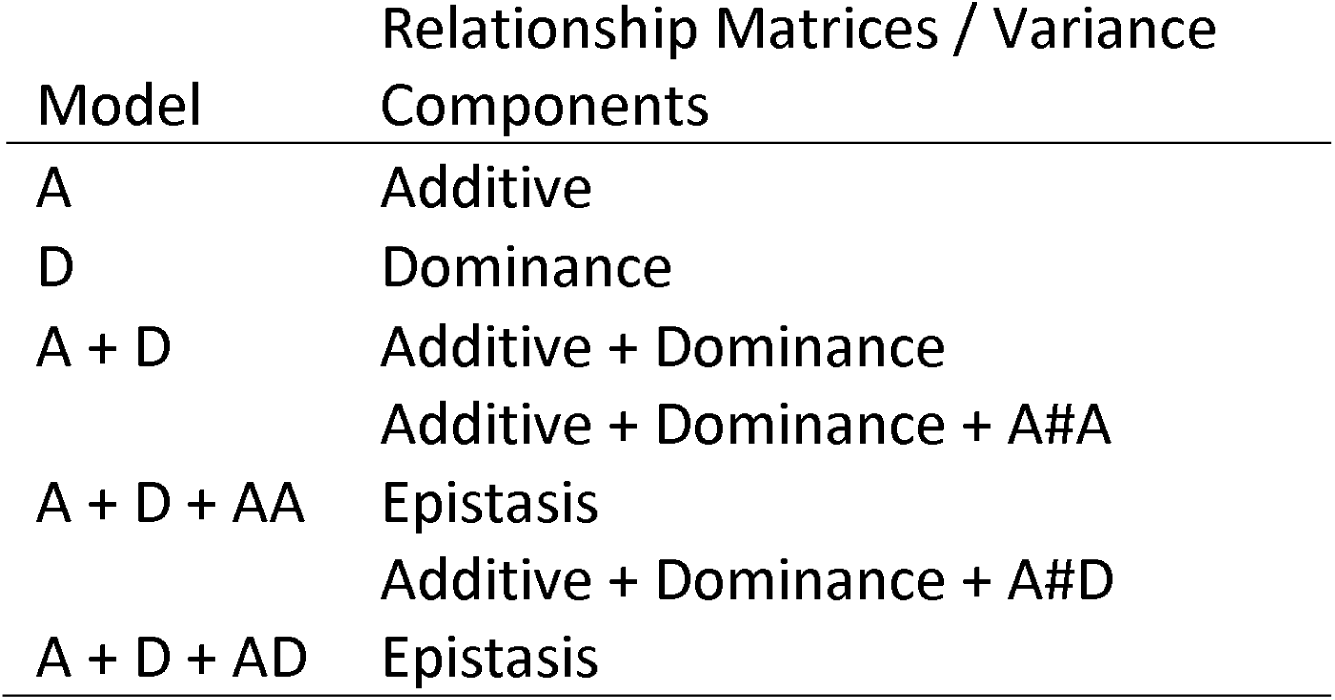
Additive plus non-additive effects models tested and their abbreviations.

| Model | Relationship Matrices / Variance Components |
| --- | --- |
| A | Additive |
| D | Dominance |
| A + D | Additive + Dominance |
|  | Additive + Dominance + A#A |
| A + D + AA | Epistasis |
|  | Additive + Dominance + A#D |
| A + D + AD | Epistasis |

The formulation described above was used to fit the GG historical data in a single model. For the C1 progenies only a single trial was available and therefore we fit all data together. Since the C1 trial was conducted across three locations but with no replications we fit the same model for C1 as GG excluding the replication term. The models described above were fit using the *regress* package in R (R Core Team 2015). The *regress* function finds REML solutions to mixed models using the Newton-Raphson algorithm.

For each trait, in both the C1 and GG we identified a “best fit” model among the models listed in Table 1 based on the lowest Akaike Information Criterion (AIC; 2*k - 2*log(likelihood), where k = number parameters estimated). We also examined the log-likelihood of each model and the total genetic variance explained (H^2^). The precision of variance component estimates and the dependency among estimates was examined using the asymptotic variance-covariance matrix of estimated parameters **(V)**. Specifically, we report standard errors for each variance component,defined as the square root of the diagonal of **V**. We also converted **V** into a correlation matrix **(F**, as in (Muñoz *et al*. 2014)), where **F** is defined as **L^-1/2^VL^-1/2^** and **L** is a diagonal matrix containing one over the square root of the diagonal of **V**. We use **F** to assess the dependency of variance components estimates, especially for comparing results among traits and populations.

#### Within-trial analyses

We used only a subset of the GG trials to estimate variance components in a single multi-environment model. In addition, we leveraged the entire historical GG data by analyzing each trial (N=47, unique location-year combinations) separately. This provided us with 47 estimates of additive, dominance and epistatic variance and we examine the distribution of variance components estimates. As in the multi-environment models, within-trial models were fit with *regress* in R.

#### Genomic prediction and cross-validation

We assessed the influence that modeling non-additive genetic variance components have on genomic prediction using a cross-validation strategy. Because single-step multi-environment models are computationally intensive, we used a two-step approach here. In the first step, we combined data from all available GG and C1 trials using the following mixed model: *y = Xβ* + *Z_rep_r* + *Z_cione_g* + *ε*. In this model, β included a fixed effect for the population mean, the location-year combination and for RTWT only, the number of plants harvested per plot. As in the single-step, multienvironment model for GG, we included the random replication effect 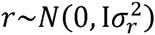. In contrast to the previous model, we did not at this stage include a genomic relationship matrix, instead we fit a random effect for clone, 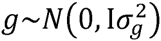, where the covariance structure was the identity matrix (I). The BLUP (ĝ) for the clone effect therefore represents an estimate of the total genetic value for each individual. The mixed model above was solved using the *lmer* function of the *lme4* R package.

In our data, the number of observations per clone ranges from one to 131 with median of two and mean of 5.97 excluding the checks TMEB1 and I30572 which had 941 and 902 observations respectively. Pooling information from multiple years and locations, especially when there is so much variation in numbers of observations can introduce bias. Much theoretical development, particularly in animal breeding has been done to address this issue, and we followed the approach recommended by Garrick et al. (Garrick *et al*. 2009). Briefly, BLUPs for clone were deregressed according to 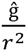 where *r*^2^ is the reliability 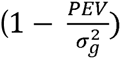 and PEV is the prediction error variances of the BLUP. In the second step of analysis, where deregressed BLUPs are used as response variables, weights are applied to the diagonal of the error variance-covariance matrix **R**. Weights are calculated as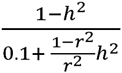, where *h*^2^ is the proportion of the total variance explained by the clonal variance component, 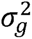 (Garrick *et al*. 2009).

We implemented a 5-fold cross-validation scheme replicated 25 times to test the accuracy of genomic prediction using the genomic relationship matrices and models described above (Table 1). In this scheme, for each replication, we randomly divided the dataset into five equally sized parts (i.e. folds). We used each fold in turn for validation by removing its phenotypes from the training population and then predicting them. We calculated accuracy as the Pearson correlation between the genomic prediction and the BLUP (g, not-deregressed) from the first step. For each model, we calculated accuracy both of the prediction from the additive kernel (where present) and the total genetic value prediction, defined as the sum of the predictions from all available kernels (e.g. additive + dominance + epistasis). Genomic predictions were made using the *EMMREML* R package.

All raw genotype and phenotype data are available on www.cassavabase.org [an exact link to be provided upon acceptance of manuscript]. Custom code used for analysis is available upon request.

## RESULTS

### Single-step, Multi-environment Variance Component Models

Our first assessments of non-additive genetic effects in cassava were single-step multi-environment models implemented in both the Genetic Gain (GG) and the Cycle 1 (C1) populations. The five models (Table 1) were initially compared using the AIC. Table 2 and 3 show the model results including AIC and variance components for the GG and C1, respectively. For HI, MCMDS, and SPROUT the best models were A + D, A + D + AD and A + D + AA, respectively. For these three traits, the best model according to AIC was the same between the GG and C1. For DM the additive only model was best in the GG but an A + D model was selected in C1. For RTWT, in the GG an A + D + AA model was selected but in the C1 the dominance only model was selected. Finally, for MCBBS the additive only model was best in the GG but A + D + AD was selected in the C1.

**Table 2.**
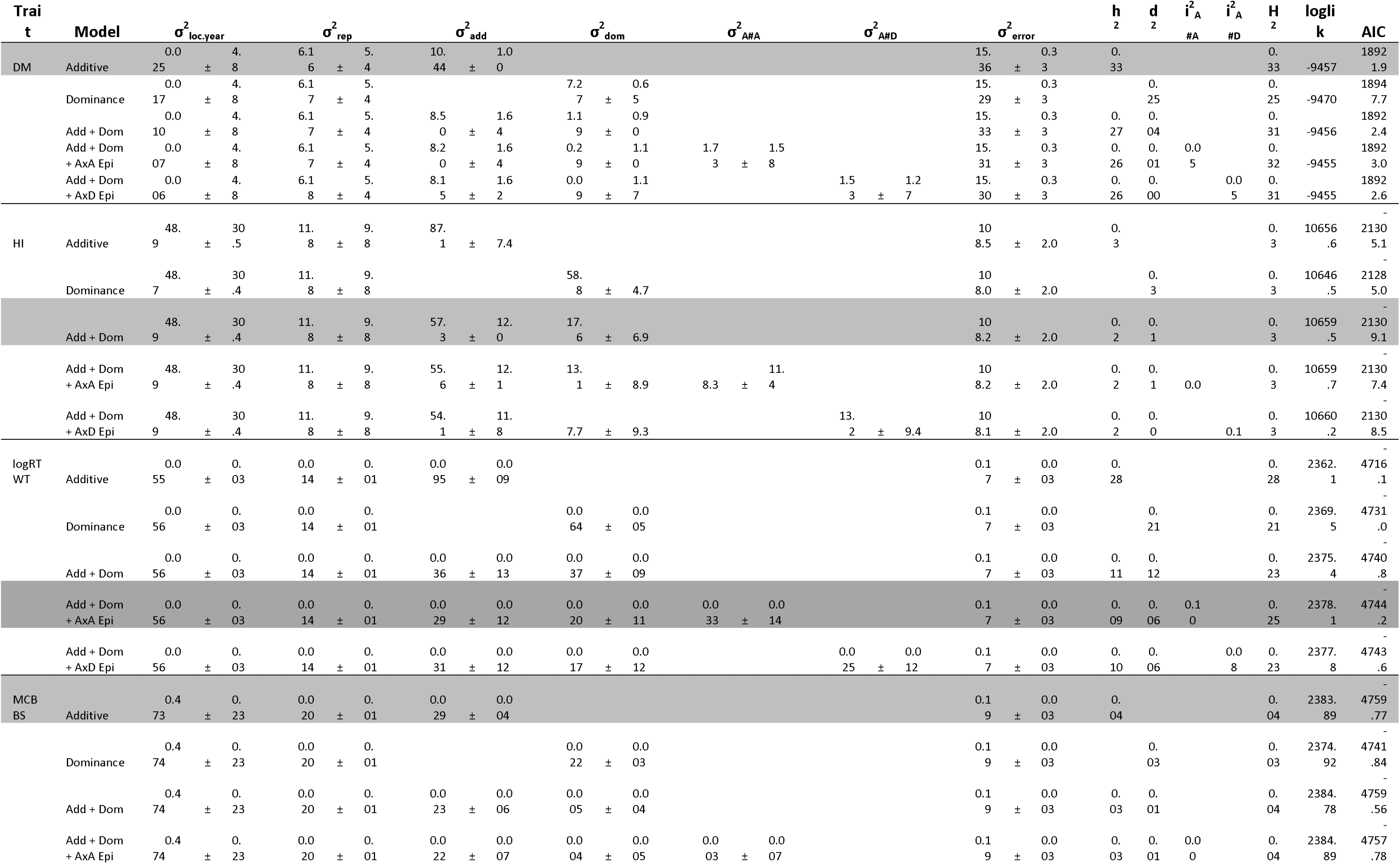
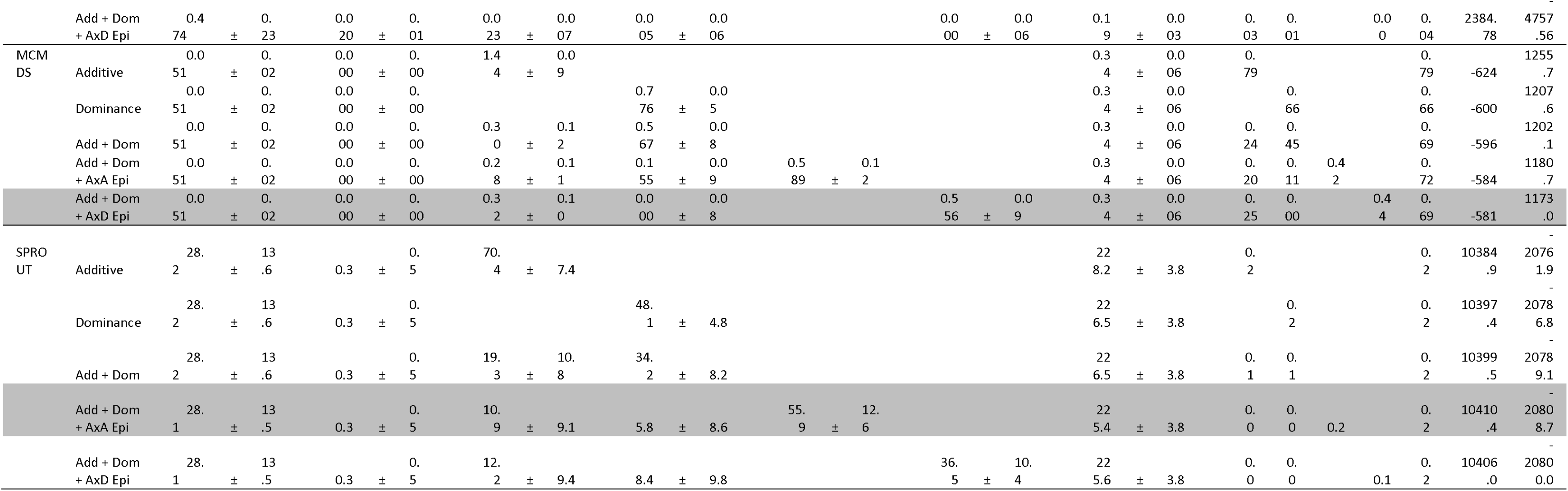
Single-step multi-environment model results for the Genetic Gain population. Variance components (± standard errors), narrow-sense heritabilities (h^2^), proportion of the total phenotypic variance explained by dominance (d^2^), additive-by-additive epistasis (i^2^_A#A_), additive-by-dominance epistasis (i^2^_A#D_) and broad-sense heritability (H^2^) are provided. Model log-likelihoods and Akaike Information Criterion (AIC) are also given. The best model for each trait (lowest AIC) is highlighted in grey.

**Table 3.**
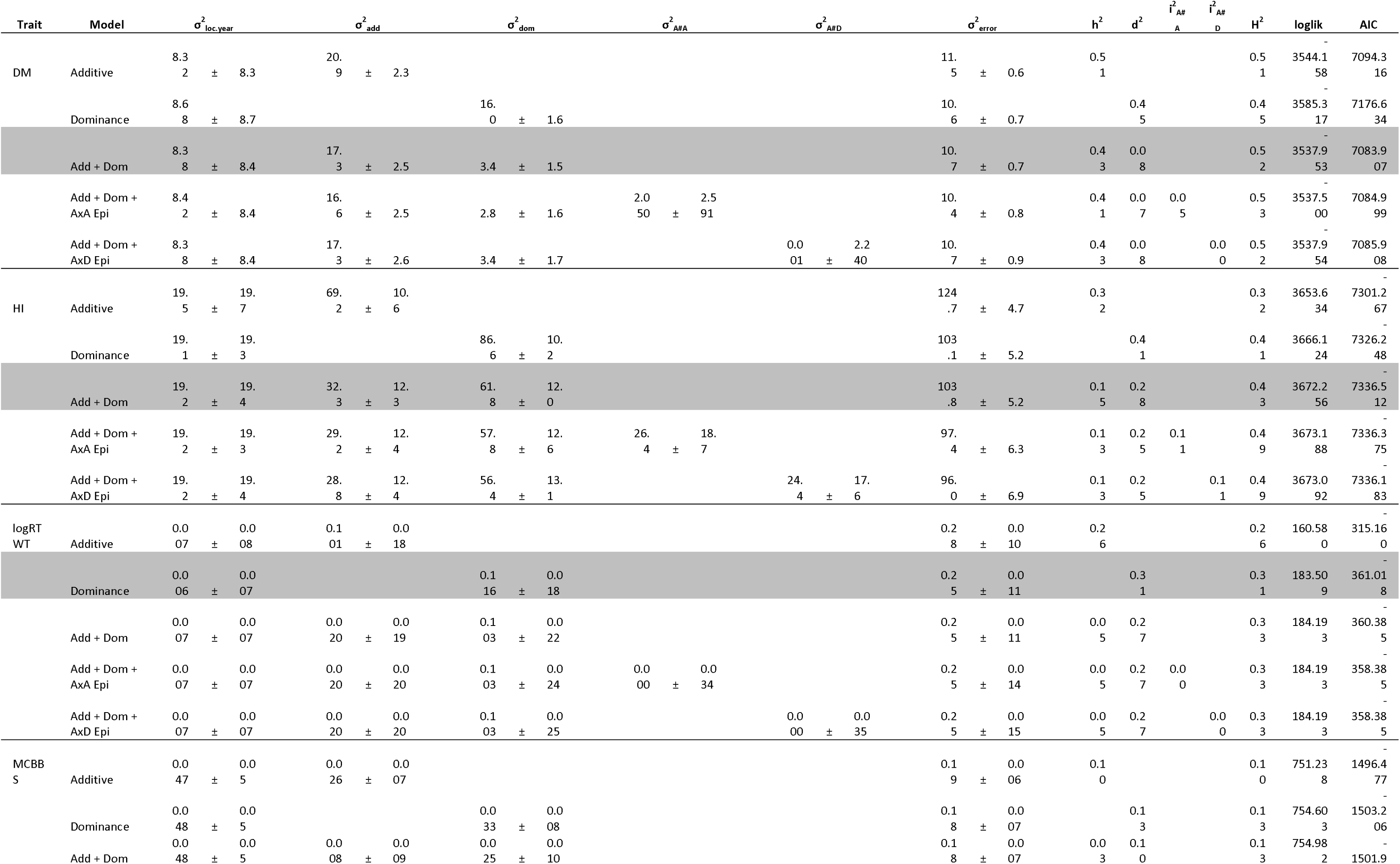
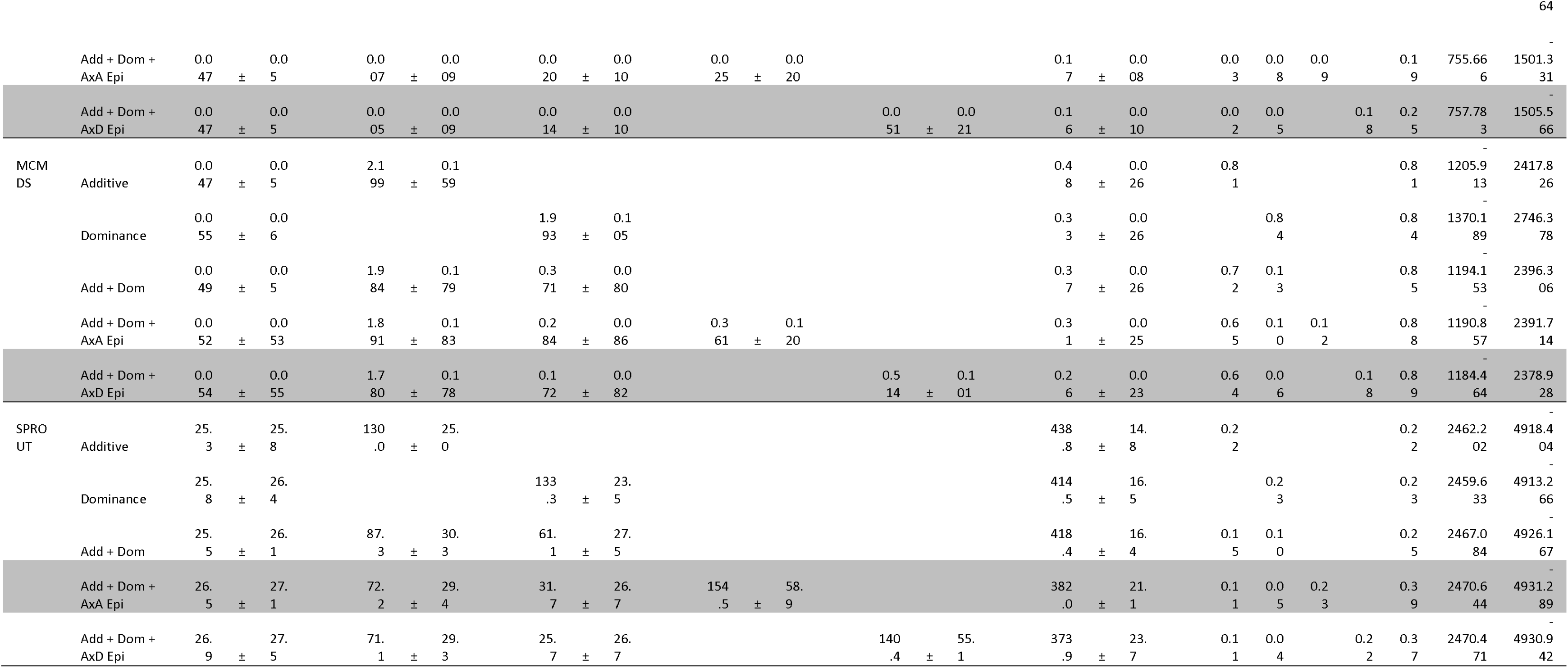
Single-step multi-environment model results for the Cycle 1 population. Variance components (± standard errors), narrow-sense heritabilities (h^2^), proportion of the total phenotypic variance explained by dominance (d^2^), additive-by-additive epistasis (i^2^_A#A_), additive-by-dominance epistasis (i^2^_A#D_) and broad-sense heritability (H^2^) are provided. Model log-likelihoods and Akaike Information Criterion (AIC) are also given. The best model for each trait (lowest AIC) is highlighted in grey.

For every trait, when comparing the model achieving the highest broad-sense heritability (H^2^), we saw that the H^2^ increased from GG to C1. This can be seen most easily in Figure 1, which shows how total explainable genetic variance (H^2^) is partitioned among variance components in the C1 and GG (also see Tables 2 & 3). In GG, the additive only model had the highest H^2^ for all traits, but in C1 additive + non-additive models always had at least slightly higher H^2^.

**Figure 1.**
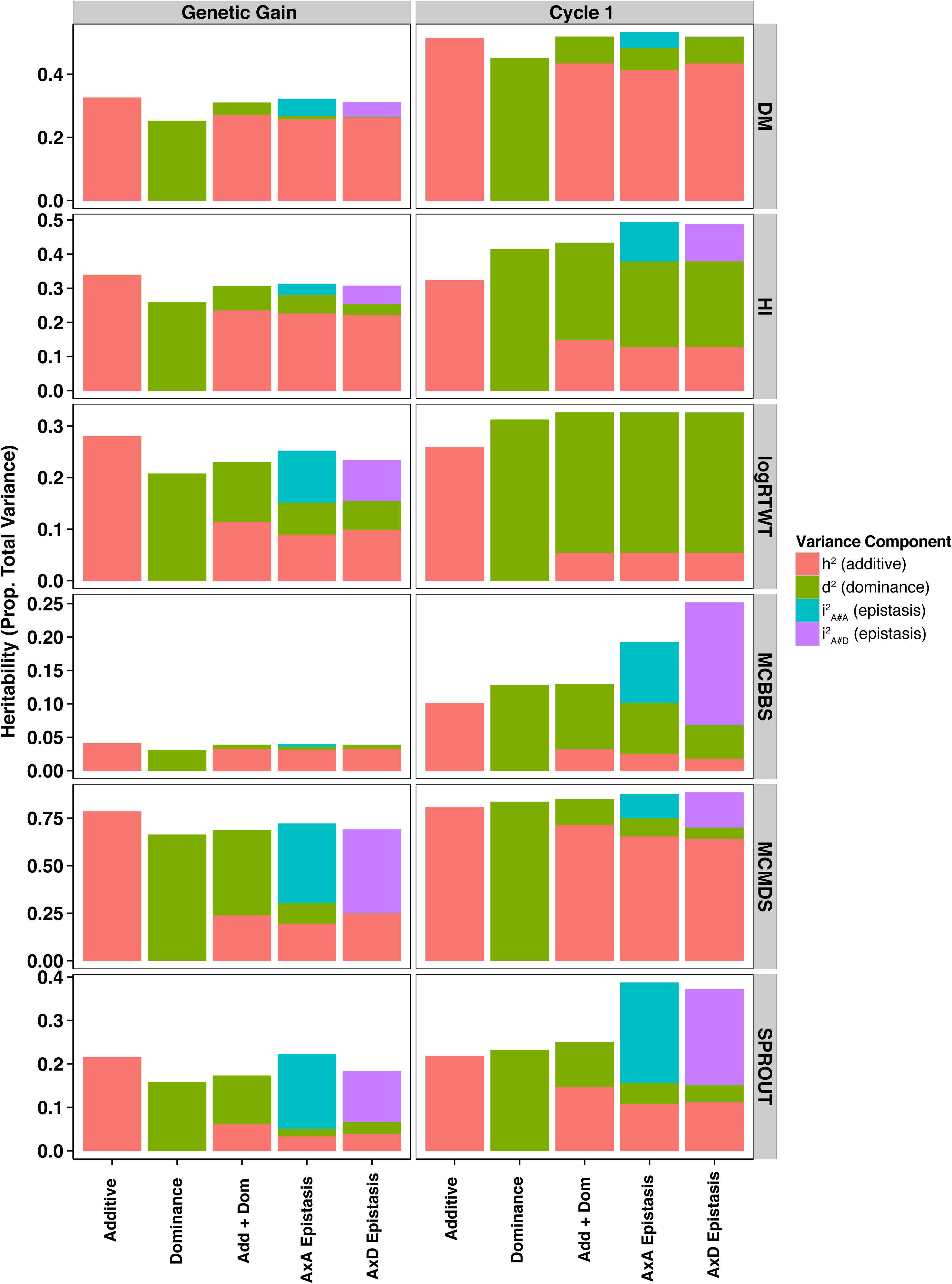
Partitioning of broad-sense heritability among variance components for single-step multi-environment models in the Genetic Gain and Cycle 1 populations for each trait. Results from each of five models are shown in each panel broken down by trait (rows) and population (columns). Models include additive only (Additive), dominance only (Dominance), Additive plus Dominance (Add_Dom), Additive plus dominance plus either AxA epistasis (Add_Add_Epistasis) or AxD epistasis (Add_Dom_Epistasis).

For DM, the additive component explained more variance across all models in the C1 compared to the GG, e.g. 0.33 (GG, Additive) and 0.51 (C1, Additive). For DM, the A + D + AA model actually had the highest H^2^ in the C1 but the variance component for A#A epistasis was not distinguishable from zero (2.05±2.59).

For HI, the additive only model was very similar between populations (h^2^ = 0.34 in GG and 0.32 in C1). But for the best model, A + D, more of the total genetic variance is partitioned to C1 (0.28) than in the GG (0.07). Like for DM, the three component models (A + D + AA and A + D + AD) explained the most variance for the C1 (H^2^ = 0.49), but the epistatic components had large standard errors (26.4±18.7 and 24.4±17.6 respectively).

RTWT was strongly non-additive in both populations. In the GG the A + D + AA model, genetics explained 25% of the phenotypic variance with non-additive variances (D + AA) explained over half of that amount (16% collectively) and the epistatic term was significantly different from zero (0.033 ± 0.014). In the C1, 31% of the phenotypic variance was explainable by dominance alone.

MCBBS had a low heritability overall (H^2^ = 0.04 in the GG for the additive only model, and 0.25 in the C1 for the A + D + AD model). While the limited apparent genetic variance in the GG was best explained additively, in the C1 92% of total genetic variance was non-additive.

For MCMDS, the additive model had the highest H^2^ (0.79) in the GG but the AIC best model, A + D + AD explained most (0.89) in the C1. The AIC best model for both C1 and GG was the same. In both cases, significant AxD epistasis was detected, explaining 44% of the variance in the GG and 18% in the C1. The additive variance increased from GG (0.25) to C1 (0.64).

SPROUT was best modeled with A + D + AA in both populations. Broad-sense heritability increased from 0.22 in GG to 0.39 in C1 for this trait. The dominance component was not significant (5.8 ± 8.6) in the GG but was in the C1 (31.7 ± 26.7).

We examined the asymptotic correlation matrices of parameter estimates (F) to ascertain the dependency of variance component estimation. Correlation matrices for every trait + model combination are provided for GG in Tables S3-S6 and for the C1 in Tables S7-S9. The correlation between genetic variance components was always negative and was, in general, higher in the GG compared to the C1. Correlations between additive and dominance components were highest in the A+D models (range -0.81 to -0.83 in the GG and -0.5 to -0.61 in the C1). Correlations between A and D components dropped in models with epistasis (range -0.42 to -0.63, GG and -0.26 to -0.58, C1). Correlations between additive and AxA epistatic variances (range -0.09 to -0.29) and AxD epistasis (range -0.07 to -0.22) were low. Correlations between dominance components and epistasis were higher ranging from -0.28 to -0.64 with AxA epistasis and -0.36 to -0.69 with AxD epistasis.

### Within-trial analyses

We also examined variance partitioning within each of 47 GG trials for the 5 models described in Table 1. The mean and variability of model parameters (variance components, heritability, etc.) across these trials are summarized in Table S10. Figure 2 provides a visual summary of the proportion of phenotypic variability explained by each genetic variance component on average across the trials. We also compared the mean AIC across trials and found them to agree overall with the results of the one-step multi-environment models (Table 2, Table S10). Specifically, the models that fit best in the one-step models were best on average in the within trial analyses for the following traits: DM (additive), HI (A+D), and MCMDS (A+D+AD). However, for RTWT (A+D instead of A+D+AD) and MCBBS (D instead of A) the within trial AIC-best models were different on average from the one-step multi-environment models.

**Figure 2.**
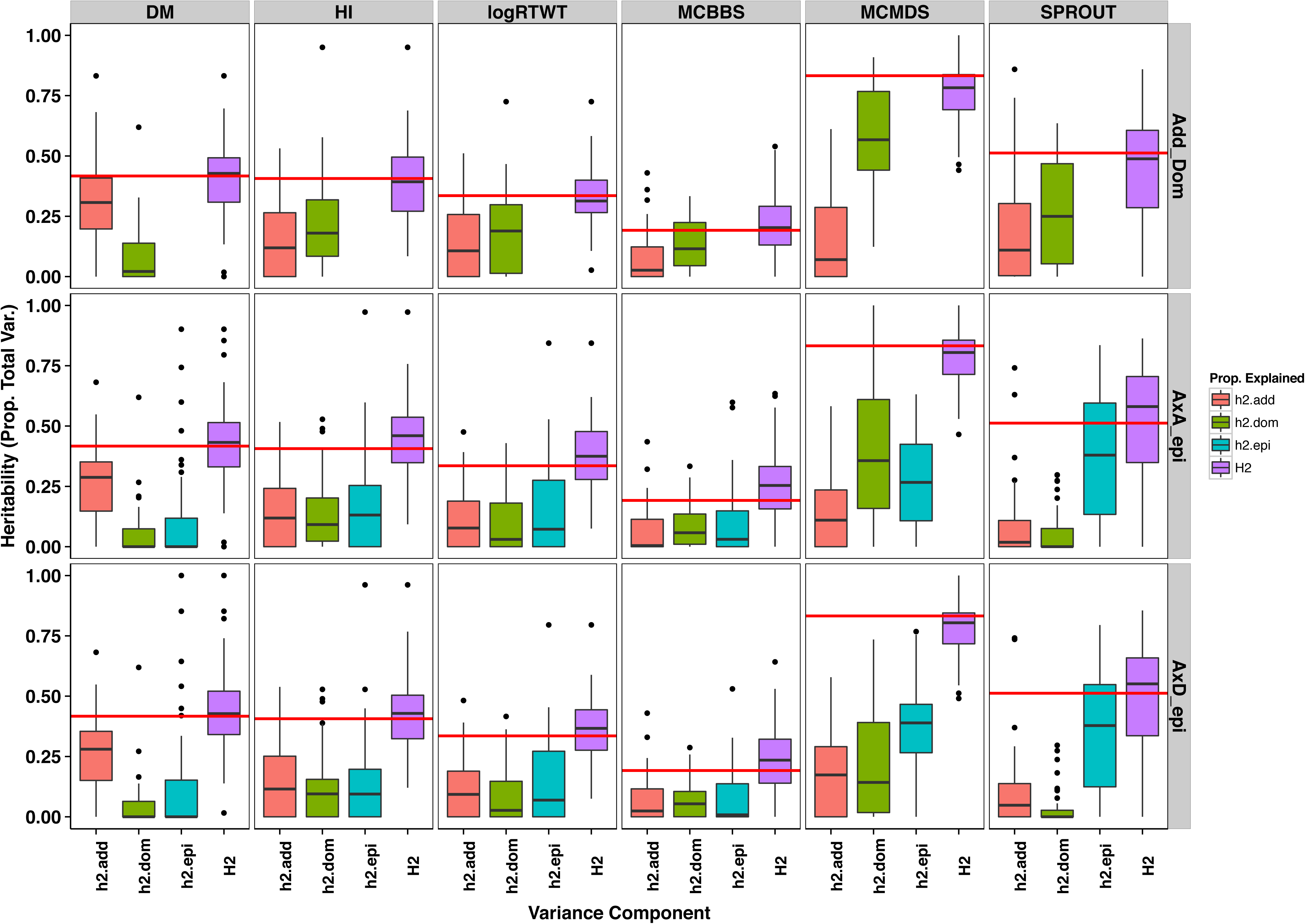
Partitioning of broad-sense heritability among variance components for the within-trial models in the Genetic Gain population. Three models were fitted for each trait in each of 47 Genetic Gain trials. Each panel contains boxplots showing the distribution proportions of the phenotypic variability explained by a corresponding variance components, including the broad-sense heritability (H^2^) across each trial. Red horizontal lines are the median narrow-sense heritability (h^2^) from the additive only model. Traits are on columns and three models are on the rows: additive plus dominance (Add_Dom), additive plus dominance plus AxA epistasis (AxA_epi) and additive plus dominance plus AxD epistasis (AxD_epi).

### Genomic Prediction of Additive and Total Genetic Value

We used cross-validation to assess the accuracy of genomic prediction of additive and total genetic value for the five models (Table 1) in both populations. For additive prediction, the additive only model had higher accuracy for every trait and every population (Figure 3, Table S11-S12). The prediction accuracy for the additive kernel was higher on average in the C1 for DM, MCBBS, MCMDS, and SPROUT compared to the GG but lower for HI and RTWT. Additive accuracies by trait (across models) were highest for DM (range: 0.54-0.60), followed by MCMDS (range: 0.44-0.54) and HI (range: 0.32-0.45). The rest of the traits had similar additive accuracies dependent on the population and model. RTWT had the overall lowest accuracy (range: 0.09-0.36).

**Figure 3.**
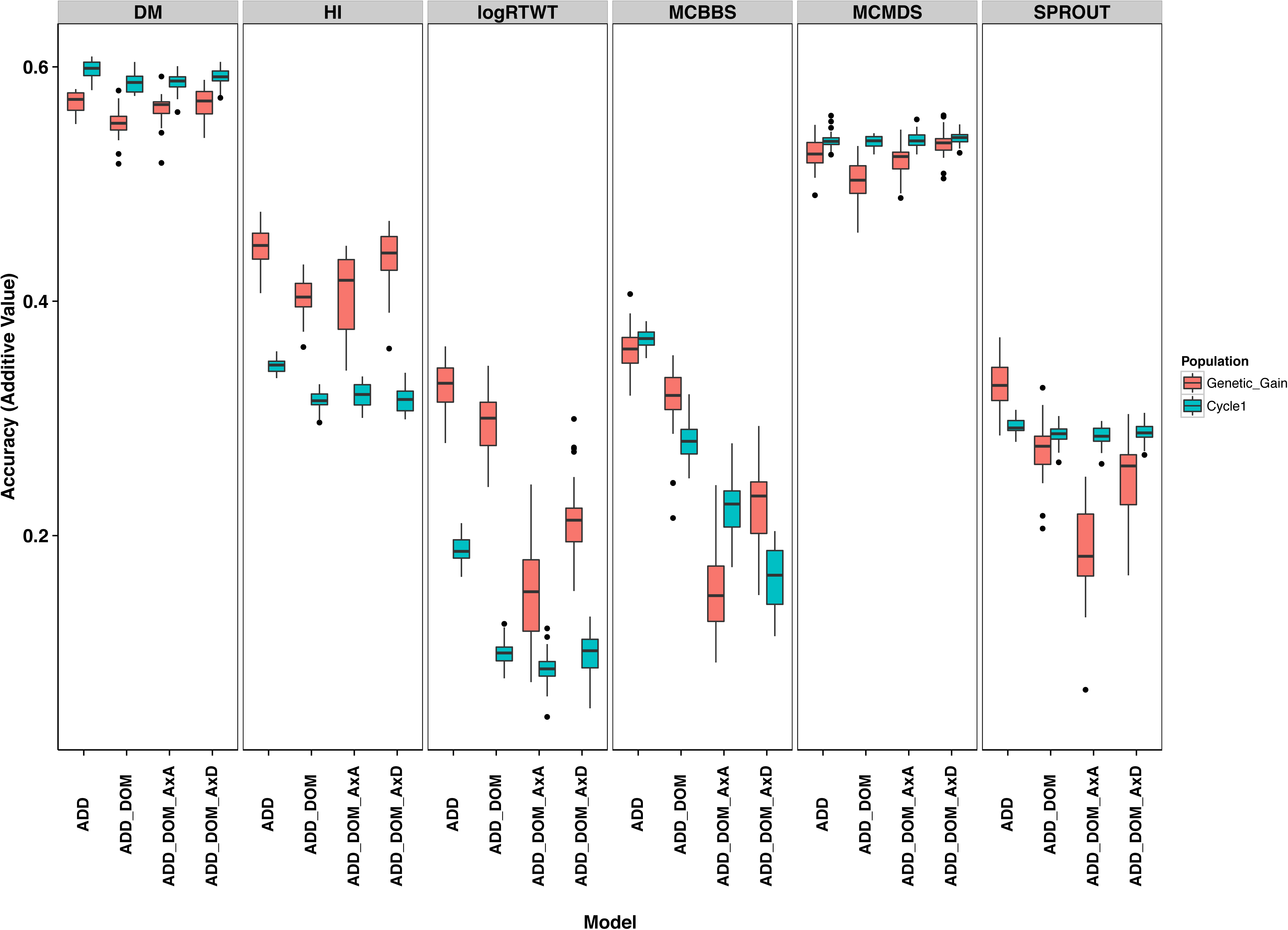
Accuracy of additive genetic value prediction in the Genetic Gain and Cycle 1 populations. Boxplots showing the distribution over 25 replicates of 5-fold cross-validation of the prediction accuracy of the additive component from four different models are shown in each panel. The accuracy within the Genetic Gain (red) and Cycle 1 (blue) are shown. Traits are in the columns. Accuracy is defined as the correlation between the prediction from the additive kernel in the model and the BLUP from the first stage of analysis where location, year and replicate variability were removed. Models included are: additive only (Add), additive plus dominance (Add_Dom), additive plus dominance plus AxA epistasis (AxA_epi) and additive plus dominance plus AxD epistasis (AxD_epi).

Across all multi-kernel analyses conducted, prediction of total genetic value was on average 42% more accurate compared to the prediction of the additive kernel in the same model. Compared to the single-kernel additive prediction however, total genetic value predictions were an average of only 6% better (maximum of 26% improvement; Figure 4, Tables S11-S12). By model, improvements in the correlation between total value and phenotype over the additive only model were 5%, 7% and 8% for A+D, A+D+AA and A+D+AD respectively. The additive only model predictions were on average 11% less accuracy in the C1 than in the GG. Additive kernel predictions from models with non-additive components were 7% lower in C1 compared to GG. Total genetic value predictions were also less accurate by 12% in the C1 relative to GG.

**Figure 4.**
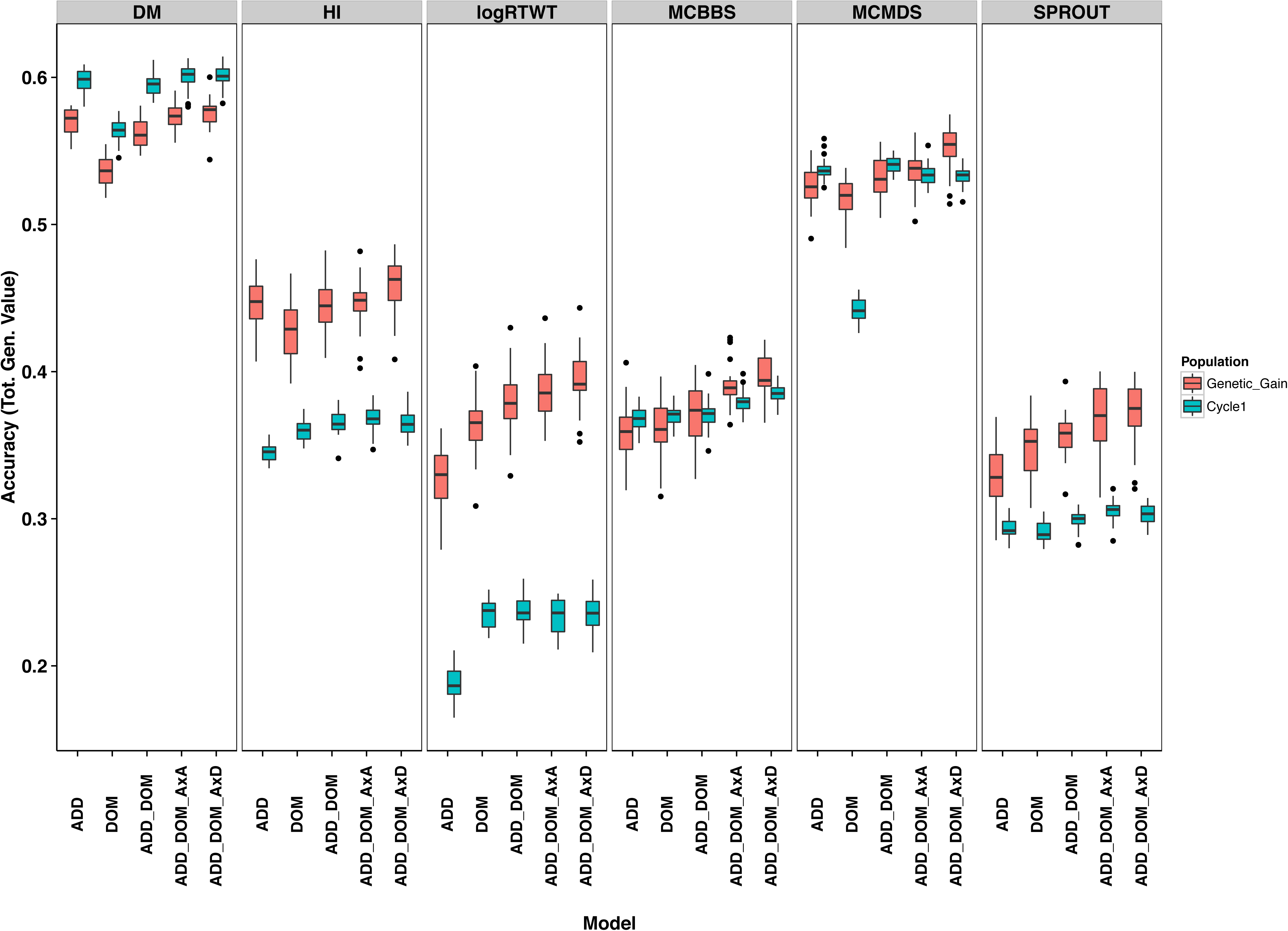
Accuracy of total genetic value prediction in the Genetic Gain and Cycle 1 populations. Boxplots showing the distribution over 25 replicates of 5-fold crossvalidation of the prediction accuracy of the total genetic value from five different models are shown in each panel. The accuracy within the Genetic Gain (red) and Cycle 1 (blue) are shown. Traits are in the columns. Accuracy is defined as the correlation between the sum of predictions from all genetic variance components in the model and the BLUP from the first stage of analysis where location, year and replicate variability were removed. Models included are: additive only (Add), dominance only (Dom), additive plus dominance (Add_Dom), additive plus dominance plus AxA epistasis (AxA_epi) and additive plus dominance plus AxD epistasis (AxD_epi).

The models we fit for multi-kernel genomic prediction involved the estimation of weight parameters corresponding to the partitioning of genetic variance among the kernels. Overall, there was a tendency for non-additive kernels (mean of 0.40 for dominance, 0.40 AxA epistasis and 0.46 AxD epistasis) to get more weight than additive (mean of 0.32). Additive prediction accuracies were positively correlated (r = 0.81) with weight placed on the additive kernel. The accuracy of total genetic value prediction was negatively correlated (r = −0.64) with weight placed on non-additive kernels (Tables S11–S12).

## DISCUSSION

In clonally propagated crops, non-additive genetic effects can be effectively exploited by the identification of superior genetic individuals as varieties. For this reason, we quantified the amount and nature of non-additive genetic variation for key traits in a genomic selection breeding population of cassava from sub-Saharan African. Then we assessed the accuracy of genomic prediction of additive compared to total (additive plus non-additive) genetic value. Using several approaches and datasets based on genome-wide marker data, we confirmed previous findings in cassava based on diallel populations, that non-additive genetic variation is significant, especially for yield traits (Cach *et al*. 2005, 2006; Calle *et al*. 2005; Jaramillo *et al*. 2005; Perez *et al*. 2005; Pérez *et al*. 2005; Zacarias and Labuschagne 2010; Kulembeka *et al*. 2012; Tumuhimbise *et al*. 2014; Ceballos *et al*. 2015; Chalwe *et al*. 2015). Further, we found that total genetic value predicted observed phenotype more accurately than additive value, although this is constrained by low broad-sense heritability and is not beneficial for traits with already high heritability (DM, MCMDS). We address the implication of these results for cassava breeding and put our work in the context of previous results in cassava, other plant and animal species below.

Our results indicate strong non-additive variance for root yields and mostly additive inheritance of root dry matter content. These findings confirm the conclusions of numerous diallel studies conducted with both Latin American (Cach *et al*. 2005, 2006; Calle *et al*. 2005; Jaramillo *et al*. 2005; Perez *et al*. 2005; Pérez *et al*. 2005) and African cassava (Zacarias and Labuschagne 2010; Kulembeka *et al*. 2012; Tumuhimbise *et al*. 2014; Chalwe *et al*. 2015) germplasm (see also (Ceballos *et al*. 2015)). In agreement with the findings of Ly et al. (Ly *et al*. 2013), we found cassava mosaic disease severity (MCMDS) to be well predicted with an additive only model. However, we found significant dominance and epistatic components in both populations analyzed. This result is in line with previous diallel studies indicating significant SCA (Tumuhimbise *et al*. 2014; Chalwe *et al*. 2015) and genetic mapping studies that identified a single major effect QTL with a dominant CMD resistance effect (Akano *et al*. 2002; Okogbenin *et al*. 2012; Rabbi *et al*. 2014). In addition, a recent genome-wide association and prediction study using non-additive genomic relationship matrices (GRMs) found that dominance and especially epistasis explain most of the variance in the region of a large-effect QTL, suggesting multiple interacting loci in the region (Wolfe et al. *in review)*. Nearly every study (but see (Zacarias and Labuschagne 2010)) indicates that Harvest Index is mostly explained by additive variance, in agreement with our findings for the GG but in contrast to our results for the C1. Our study is the first to report on the modes of inheritance of CBB and SPROUT.

The importance of non-additive genetic variance in evolution by natural and artificial selection is controversial (Hill *et al*. 2008; Crow 2010; Hansen 2013). Nevertheless, numerous studies have found and exploited dominance and epistasis in animal breeding, including dairy (Ahlborn-Breier and Hohenboken 1991; Fuerst and Sölkner 1994; Varona *et al*. 1998; Van Tassell *et al*. 2000; Palucci *et al*. 2007) and beef (Rodriguezalmeida *et al*. 1995) cattle. Diallel studies have indicated significant SCA for maize grain yield (Doerksen *et al*. 2003; Wardyn *et al*. 2007). Aside from cassava, breeding of other non-inbred, clonally propagated species also identify and make use of non-additive effects, including potato (Killick 1977), Eucalyptus (Costa E Silva *et al*. 2004) and loblolly pine (Muñoz *et al*. 2014). More recently, marker-based and GRM-based models have identified significant non-additive effects in pigs (Su *et al*. 2012; Nishio and Satoh 2014), mice (Vitezica *et al*. 2013), beef cattle (Bolormaa *et al*. 2015), dairy cows (Morota *et al*. 2014), maize (Dudley and Johnson 2009), soy (Hu *et al*. 2011), loblolly pine (Muñoz *et al*. 2014) and apple (Satish Kumar, Claire Molloy, Patricio Muñoz, Hans Daetwyler, David Chagné 2015). Results from the present study suggest that the combination of genomic selection and hybrid breeding strategies should increase the rate of genetic gain for complex traits such as yield. However, initial investment in identification of complementary heterotic groups with good specific combining ability is required.

One of the more interesting aspects of our study relative to previous ones is the comparison between a parental generation (the Genetic Gain) and their offspring (Cycle 1), a collection of full- and half-sib families. From GG to C1, the H^2^ generally increased. For RTWT, MCBBS and HI this is largely attributable to increased non-additive variance and CMD for which non-additive variance dropped in C1 relative to GG. In contrast to our result, theory suggests that reduction (or fixation) of allele frequencies at some loci relative to others in populations undergoing bottlenecks (Goodnight 1988), inbreeding (Turelli and Barton 2006) or truncation selection (Hallander and Waldmann 2007) should cause a conversion of non-additive (where present) to additive variance. These results have, however, been based on models with finite numbers of loci in linkage equilibrium. Based on the mean diagonal of the kinship matrix, C1 (0.66) does not appear notably more inbred than GG (0.64). We also calculated mean pairwise LD (GG = 0.27, C1 = 0.29) and mean LD block size (21.7 kb in GG and 23.1 kb in C1) using PLINK (version 1.9, https://www.cog-genomics.org/plink2) and found the two generations to be similar.

Probably the strongest explanation for the difference in genetic variance components between GG and C1 is the family structure (137 full-sib families from 83 outbred parents). In a population of half-sibs ¾ of the dominance variance is expressed within families and all of it for full-sib populations (Hallauer *et al*. 2010; Ceballos *et al*. 2015). Indeed, increasing the number of full-sib relationships is known to increase the non-additive genetic variance detectable in a population (Varona *et al*. 1998; Van Tassel *et al*. 2010).

It is also conceivable that maternal plant effects could increase apparent non-additive effects in C1. The C1 clones in contrast to the GG clones are new, and were derived from stem cuttings of seedling plants germinated in the previous field season (2012-2013). The suggestion is therefore that the quality and vigor of the seedling plant, giving rise to the C1 clones may influence their performance in the 2013-2014 trial. We further caution that comparison of GG and C1 may be biased by the disproportionate amount of data available for the GG.

In our study, when additive and non-additive kernels were used together, the ability of the additive kernel to predict the phenotype decreased, suggesting that the additive kernel by itself was absorbing non-additive variance. Estimates of additive genetic variance have previously been shown to capture some non-additive effects (Lu *et al*. 1999; Su *et al*. 2012; Zuk *et al*. 2012; Muñoz *et al*. 2014). Predictions models that do not explicitly incorporate non-additive components may therefore achieve gains in the short-term that break down over the long-term (Cockerham and Tachida 1988; Walsh 2005; Hansen 2013). Including non-additive GRMs when estimating additive genetic (i.e. breeding) values may therefore provide a less biased, more accurate selection of parents for genomic selection (Costa E Silva *et al*. 2004; Palucci *et al*. 2007). We note that our predictions of total genetic value were focused on parametric models based in quantitative genetic theory. However, many non-parametric and non-linear approaches are available (e.g. RKHS and random forests) that may capture even more non-additive variation than found in our study.

Non-additive variation is prevalent in cassava, especially for low heritability traits. This has many important implications for cassava breeding. It explains, in part, why genetic gains have been slow (Ceballos *et al*. 2012). Inbreeding to convert dominance variance to additive and better control epistatic combinations, as in maize, has been suggested as a solution to non-additive genetics (Ceballos *et al*. 2015). Even for low h^2^ traits and without inbred cassava, using the kinds of models presented in this paper, good parents can be selected based on additive predictions and total genetic value can be simultaneously estimated for the identification of potential commercial varieties, all based on the combination of marker and preliminary field trial data(Heslot and Mark 2015). This approach has been previously advocated for plant breeding (Oakey *et al*. 2007; Heslot and Mark 2015) and has proven effective in animal breeding, e.g. (Ahlborn-Breier and Hohenboken 1991; Palucci *et al*. 2007; Su *et al*. 2012; Nishio and Satoh 2014). Non-additive models using genomic relationship matrices can thus improve the efficiency and productivity of variety selection pipelines that are the most labor and time intensive part of selecting good cassava clones after crossing.

## Acknowledgements

We acknowledge the Bill & Melinda Gates Foundation and UKaid (Grant 1048542; http://www.gatesfoundation.org) and support from the CGIAR Research Program on Roots, Tubers and Bananas (http://www.rtb.cgiar.org). We give special thanks to A. G. O. Dixon for his development of many of the breeding lines and historical data we analyzed. Thanks also to A. I. Smith and technical teams at IITA for collection of phenotypic data and to A. Agbona and P. Peteti for data curation.

---

**Supplementary Table 1.** Pedigree and related information for the IITA: Genetic Gain population, Nigeria.

**Supplementary Table 2.** Sample size (Nobs), replication number (Nreps), clone number (Nclones) and whether a trial (LOC.YEAR) was included in one-step multi-environment models for both populations analyzed.

**Supplementary Table 3.** Pedigree information for the IITA: GS Cycle 1 population, Nigeria.

**Supplementary Table 4.** Asymptotic correlation matrices of parameter estimates representing the dependency of variance component estimation for the additive plus dominance model in the Genetic Gain population, by trait.

**Supplementary Table 5.** Asymptotic correlation matrices of parameter estimates representing the dependency of variance component estimation for the additive plus dominance plus additive-by-additive epistasis model in the Genetic Gain population, by trait.

**Supplementary Table 6.** Asymptotic correlation matrices of parameter estimates representing the dependency of variance component estimation for the additive plus dominance plus additive-by-dominance epistasis model in the Genetic Gain population, by trait.

**Supplementary Table 7.** Asymptotic correlation matrices of parameter estimates representing the dependency of variance component estimation for the additive plus dominance model in the Cycle 1 population, by trait.

**Supplementary Table 8.** Asymptotic correlation matrices of parameter estimates representing the dependency of variance component estimation for the additive plus dominance plus additive-by-additive epistasis model in the Cycle 1 population, by trait.

**Supplementary Table 9.** Asymptotic correlation matrices of parameter estimates representing the dependency of variance component estimation for the additive plus dominance plus additive-by-dominance epistasis model in the Cycle 1 population, by trait.

**Supplementary Table 10.** Within-trial model results for the Genetic Gain population. The mean (± standard errors) across 47 trials for each trait and model fitted is given for the following model parameters: variance components, narrow-sense heritabilities (h^2^), proportion of the total phenotypic variance explained by dominance (d^2^), additive-by-additive epistasis (i^2^_A#A_), additive-by-dominance epistasis (i^2^_A#D_) and broad-sense heritability (H^2^), trial sample size (N), model log-likelihoods and Akaike Information Criterion (AIC) are also given. The best model for each trait (lowest AIC) is highlighted in grey.

**Supplementary Table 11.** Results from 25 replicates of 5-fold cross-validation in the Genetic Gain population are given. Mean (± standard errors) across the 25 replicates are given for prediction accuracy of each kernel plus total genetic value (sum across all kernels), variance components (V_g_ and V_e_) and kernel weights.

**Supplementary Table 12.** Results from 25 replicates of 5-fold cross-validation in the Cycle 1 population are given. Mean (± standard errors) across the 25 replicates are given for prediction accuracy of each kernel plus total genetic value (sum across all kernels), variance components (Vg and Ve) and kernel weights.

